# A CD24^+^CD271^+^ Melanoma Cancer Stem Cell Possesses Hybrid Characteristics of its Single Marker Counterparts and Promotes Invasion and Therapeutic Resistance

**DOI:** 10.1101/2023.06.07.544036

**Authors:** Olivia Knowles, Patricio Doldan, Isabella Hillier-Richardson, Stephanie Lunt, Gehad Youssef, Luke Gammon, Ian C. Mackenzie, Michael P. Philpott, Hasan Rizvi, Daniele Bergamaschi, Catherine A. Harwood, Adrian Biddle

**Author notes:** Corresponding author: Adrian Biddle, Centre for Cell Biology and Cutaneous Research, Blizard Institute, 4 Newark Street, London, E1 2AT, UK. These authors contributed equally (Olivia Knowles, Patricio Doldan, Isabella Hillier-Richardson). Significance: We identify a stem cell hierarchy in melanoma using a novel combination of cell surface markers, and demonstrate a surprising confluence of the attributes of two discrete cellular sub-populations into a hybrid population with enhanced stem cell characteristics. The CD24^+^ and hybrid CD24^+^CD271^+^ melanoma stem cells are resistant to targeted melanoma therapeutic agents and are present in a subset of human melanomas.

## Abstract

An important role for phenotype switching has been demonstrated in metastasis and therapeutic resistance of both melanoma and epithelial tumours. Phenotype switching in epithelial tumours is driven by a minority cancer stem cell sub-population with lineage plasticity, but such a sub-population has not been identified in melanoma. We investigated whether cell surface markers used to identify cancer stem cells in epithelial tumours could identify a cancer stem cell sub-population with lineage plasticity in melanoma. We identified a CD24^+^CD271^+^ minority sub-population in melanoma that possesses the stem cell characteristics of lineage plasticity and self-renewal. This population displayed hybrid characteristics, combining the attributes of discrete CD24^+^ and CD271^+^ cellular sub-populations but with heightened sphere formation, lineage plasticity, migratory ability and drug resistance over its single-marker counterparts. CD24^+^ and CD24^+^CD271^+^ stem cell sub-populations were observed in 10% of human melanomas, mainly at the invasive front. The lack of CD24^+^ and CD24^+^CD271^+^ stem cells in the majority of human melanoma specimens led us to conclude that they may be dispensable for melanoma progression. Nevertheless, the enhanced sphere formation, lineage plasticity, migratory ability and drug resistance of the CD24^+^CD271^+^ sub-population may signal a contextual requirement for these stem cells when melanomas face challenging environments both clinically and in experimental systems.

## Introduction

Melanoma is a highly aggressive form of skin cancer. Invasive melanoma accounts for about 1% of all skin cancers, but is responsible for around 60% of skin cancer deaths (American Cancer Society.). Melanoma is unusual in that it is often very invasive at an early stage of tumour development, setting it apart from other solid tumours that typically become invasive at a later stage. This single attribute is a major contributor to melanoma’s high mortality rate, despite recent therapeutic advances. Even when a patient presents with a small primary lesion, the tumour has often already invaded beyond the primary site and is thus incurable with surgery. Melanoma develops from melanocytes, the pigment forming cells of the skin. These are distinct from the other cells of the skin in that they are not derived from an epithelial developmental lineage. Instead, they derive from the neural crest, a lineage that undergoes an epithelial-mesenchymal-transition (EMT) and migrates throughout the embryo to produce a number of different cell types in development (Morrison et al., 1999).

EMT is a major driver of tumour invasion in epithelial cancers (Biddle et al., 2011, Lu and Kang, 2019), with activation of an EMT program during tumour development greatly enhancing metastatic spread (Tsai et al., 2012, Pastushenko et al., 2021). Activation of an EMT program is restricted to cancer stem cells (CSCs), a minority sub-population of cells within the tumour that possess tumour-initiating potential by virtue of their self-renewal and lineage plasticity. Previous work in epithelial cancers has shown that tumour invasion and metastasis is driven by phenotypic switching between two CSC states: 1) a stationary, highly proliferative, drug-sensitive epithelial state that is CD44^high^, EpCAM^high^ and Vimentin^low^, and 2) an invasive, slow-growing, drug-tolerant mesenchymal-like state that is CD44^high^, EpCAM^low^ and Vimentin^high^ (Biddle et al., 2016, Biddle et al., 2011, Liu et al., 2014, Hermann et al., 2007). There is further heterogeneity within the sub-population of cells that have transitioned through EMT into a mesenchymal-like state, with only some of these cells retaining the CSC attribute of lineage plasticity that enables them to regenerate a proliferative epithelial tumour at a metastatic site and thus propagate metastatic dissemination (Pastushenko et al., 2018, Biddle et al., 2011). Possessing the two key CSC attributes of self-renewal and lineage plasticity, we have shown in oral squamous cell carcinoma (OSCC) that these metastatic EMT-CSCs can be identified by their expression of the plasticity marker CD24 alongside the epithelial marker EpCAM, the mesenchymal marker Vimentin, and the CSC marker CD44 (Youssef et al., 2023, Biddle et al., 2016).

As melanomas derive from a lineage that has already undergone a developmental EMT, they do not need to undergo the phenotypic switching described above. It might be expected that this is a key underlying reason for their propensity for early tumour invasion. Indeed, melanoma cells do not transition between epithelial and mesenchymal states, and ubiquitously express high levels of the mesenchymal marker Vimentin (Rambow et al., 2019). However, numerous studies have demonstrated that melanomas nevertheless exhibit lineage plasticity and transition between two states that are analogous to those seen in epithelial tumours: 1) a stationary, highly proliferative, drug-sensitive melanocytic state that is MITF^high^, Axl^low^, Zeb1^low^ and CD271^low^, and 2) an invasive, slow-growing, drug-tolerant neural crest-like state that is MITF^low^, Axl^high^, Zeb1^high^ and CD271^high^ (Carreira et al., 2006, Richard et al., 2016, Müller et al., 2014, Restivo et al., 2017, Sensi et al., 2011). This demonstrates the same important role for lineage plasticity as has been seen in epithelial tumours.

We therefore hypothesised that the minority CSC sub-population retaining lineage plasticity in epithelial tumours may have a parallel state in melanoma. In order to investigate this, we incorporated a previously reported melanoma CSC marker, CD271 (Boiko et al., 2010), alongside the markers we previously identified in epithelial carcinomas. As well as being a reported melanoma CSC marker, CD271 is a key cell surface marker for the invasive slow-growing state in melanoma (Restivo et al., 2017). It is also a marker for resistance to BRAF inhibitors (Richard et al., 2016, Patel et al., 2020) and resistance to immunotherapy (Landsberg et al., 2012). We therefore selected CD271 for investigation alongside the cell surface markers for plastic CSCs in epithelial tumours - CD24, EpCAM and CD44 (Youssef et al., 2023, Biddle et al., 2016). We sought to determine whether any of these markers in combination could discriminate a CSC sub-population in melanoma.

## Results

We initially cultured 4 established human melanoma cell lines – LM4, LM32, CHL-1 and A375M. We assessed each cell line for cell surface expression of CD44, EpCAM, CD24 and CD271 by flow cytometry (Figure 1A). All cell lines were homogeneously CD44^+^EpCAM^-^, except for CHL-1 which contained a small number of CD44^-^EpCAM^-^ cells. This demonstrates the mesenchymal character of all cell lines, in agreement with the ubiquitously high expression of the mesenchymal marker Vimentin in melanoma (Rambow et al., 2019). It also suggests that CD44, a CSC marker in epithelial cancers, does not discriminate sub-populations in melanoma. The surface expression of CD24 and CD271 was more variable between cell lines; the 4 lines expressed varying levels of CD271, whereas only CHL-1 expressed CD24. This gave rise to both CD24^+^CD271^+^ and CD24^+^CD271^-^ sub-populations in the CHL-1 cell line only. Brightfield imaging (Figure 1B) demonstrated a mesenchymal morphology for all cell lines, further underlining their mesenchymal character. However, there was heterogeneity between cell lines, with LM4 and LM32 having a homogenously mesenchymal morphology whereas CHL-1 and A375M were more heterogeneous in their morphology, with clustering cobblestone regions. These were more pronounced in CHL-1, which overall showed a greater level of morphological heterogeneity than the other three lines. In order to assess the CSC attributes of the four lines, we performed a tumoursphere assay. The tumoursphere assay tests ability to survive and grow in suspension, and is a surrogate assay for CSC identity (Mukherjee et al., 2021); it enriches for cells that exhibit resistance to the oxidative stress and anoikis triggered by suspension culture, and continue to self-renew under these conditions (Zhu et al., 2020). We performed the tumoursphere assay under two conditions; in fully supplemented FAD medium with 10% serum, as used to maintain a proliferative undifferentiated phenotype in adherent culture (Figure 1C, top), or in a defined serum-free medium that was developed for organoid culture and provokes differentiation (Figure 1C, bottom) (Sato et al., 2011). Interestingly, behaviour was starkly different under the two conditions; all four cell lines produced both primary and secondary spheres in the FAD medium, whereas only CHL-1 produced spheres in the more stringent serum-free conditions. Therefore, only the CHL-1 cell line retains a self-renewing CSC population under differentiation culture conditions.

**Figure 1.**
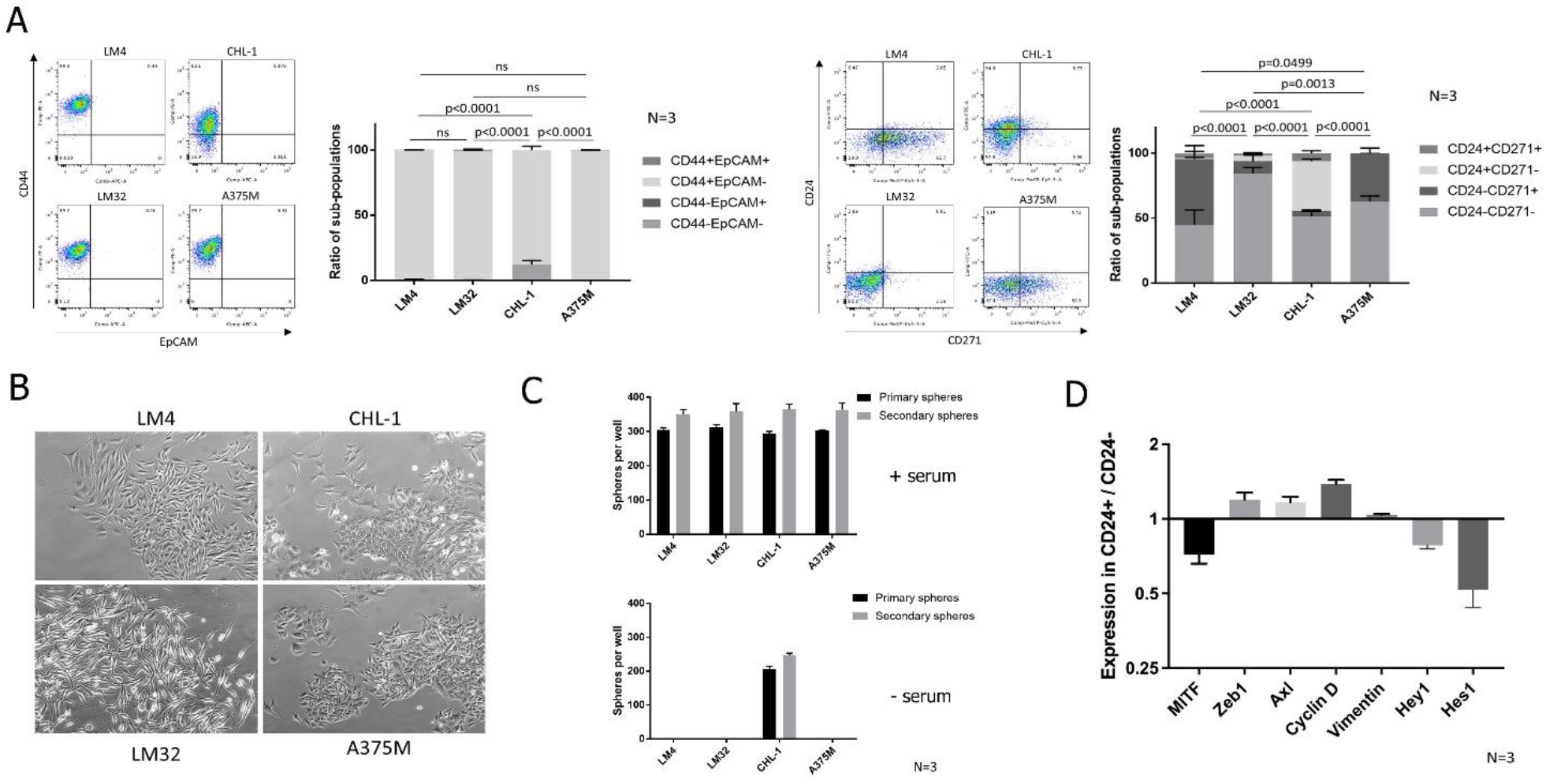
Cellular heterogeneity and stem cell attributes differ between melanoma cell lines. **A**, Flow cytometric analysis of the four cell lines for CD44 and EpCAM (left) and CD24 and CD271 (right). The graphs show mean +/-SEM of sub-populations in each of 4 quadrants based on +/-gating set using isotype controls, as a percentage of the total cells, presented as stacked bars. Representative flow cytometry plots for each cell line are to the left of each graph. **B**, Brightfield images of four melanoma cell lines in culture. **C**, Sphere counts for the four cell lines, for primary spheres and secondary spheres after dissociation and replating, in serum-containing FAD medium (top) and serum-free medium (bottom). **D**, Gene expression in FACS sorted CD24+ vs CD24-sub-populations. For each gene, expression was measured by QPCR and is represented as the expression level in CD24+ cells relative to that in CD24-cells. For each experiment, the number of biological repeats is indicated next to the graph. P-values were calculated using two-way ANOVA.

Thus, of the four melanoma cell lines, only CHL-1 exhibits heterogeneous sub-populations based on any of the CSC markers derived from epithelial tumours. Both CD44 and CD24 were heterogeneously expressed in CHL-1, but only CD24 identified a minority positive sub-population that was absent from the other cell lines. CHL-1 was also the only cell line capable of producing tumourspheres under the more stringent serum-free conditions. We therefore focussed on CD24 as a potential marker of a sphere-forming CSC sub-population in CHL-1.

We performed QPCR analysis on FACS sorted CD24^+^ and CD24^-^ sub-populations from CHL-1, for markers of the two recognised melanoma phenotypic states – melanocytic (proliferative) and neural crest-like (invasive) (Figure 1D). Compared to CD24^-^ cells, CD24^+^ cells had heightened expression of markers of the neural crest-like state (Zeb1 and Axl) and reduced expression of the melanocytic state marker MITF. The mesenchymal marker Vimentin was stably expressed across the two sub-populations. Interestingly, and contrary to the documented growth-arrested status of the neural crest-like state, the CD24^+^ sub-population had greatly increased Cyclin D expression. This suggested that CD24 may mark an alternative phenotypic state that has a dedifferentiated neural crest-like status whilst maintaining proliferative competence, as has been recently proposed (Rambow et al., 2019). The Notch signalling effectors Hes and Hey were reduced in the CD24^+^ sub-population, which is contrary to the documented role of Notch signalling in the CD271^+^ neural crest-like state (Murtas et al., 2017, Filipp et al., 2019). This is further evidence that CD24 may mark an alternative phenotypic state. Therefore, we focussed on CD24 alongside CD271 for further investigations in the CHL-1 cell line in order to determine whether this cell line contains a CSC sub-population that is absent from the other melanoma cell lines.

We FACS sorted four sub-populations (CD24^+^CD271^+^, CD24^+^CD271^-^, CD271^+^CD24^-^ and CD24^-^CD271^-^) from the CHL-1 sub-line and performed a range of functional CSC assays. Using the double-negative CD24^-^CD271^-^ sub-population as a baseline, we found that only the CD24^+^CD271^+^ sub-population produced a significantly increased number of tumourspheres (using the more stringent serum-free condition) (Figure 2A). It also produced an increased number of colonies in a colony formation assay, another test for self-renewal (Figure 2B). Therefore, the CD24^+^CD271^+^ sub-population exhibits enhanced anoikis resistance and self-renewal. To assess lineage plasticity, we cultured each sorted sub-population for 7 days and then re-assessed the sub-population distribution (Figure 2C). Plasticity was much more pronounced in the CD24^+^CD271^+^ sub-population, which gave rise to large numbers of cells in each of the other three sub-populations. In contrast, the CD24^+^CD271^-^ and CD24^-^CD271^+^ sub-populations gave rise mostly to CD24^-^CD271^-^ cells, and the CD24^-^CD271^-^ sub-population produced few cells in any of the other sub-populations. Therefore, the CSC attributes of self-renewal and lineage plasticity reside mainly in the CD24^+^CD271^+^ sub-population.

**Figure 2.**
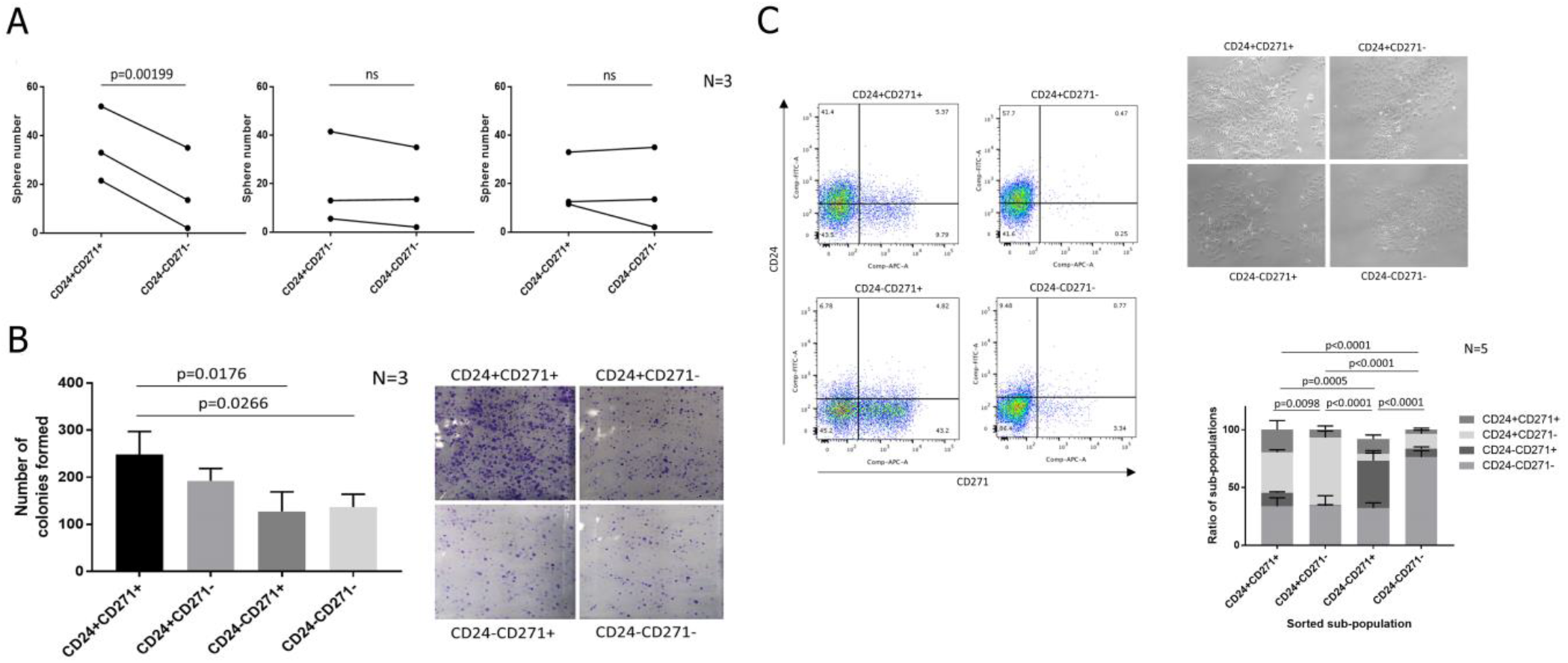
CD24 and CD271 mark a stem cell sub-population in the CHL-1 melanoma line. **A**, Primary sphere counts for four FACS sorted sub-populations from the 4 quadrants based on +/-gating set using isotype controls. Results are presented as paired individual data points (due to considerable baseline variation between repeats), with each of the three marker-positive sub-populations compared individually to the CD24-CD271-sub-population. P-values were calculated using paired ANOVA (due to the variation between repeats). **B**, Colony counts in 2D culture for the four FACS sorted sub-populations, with representative images of crystal violet stained colonies for each sub-population. P-values were calculated using unpaired ANOVA. **C**, Flow cytometric analysis for CD24 and CD271, 7 days after sorting and re-plating the four sub-populations, with accompanying brightfield images of the four sorted sub-populations in culture. The graph shows mean +/-SEM of sub-populations in each of 4 quadrants based on +/-gating set using isotype controls, as a percentage of the total cells, presented as stacked bars. P-values were calculated using two-way ANOVA. Representative flow cytometry plots for each re-plated sub-population are to the left of the graph. For each experiment, the number of biological repeats is indicated next to the graph.

We next assessed the ability of the four CHL-1 sub-populations to drive invasive dissemination. The CD24^+^CD271^+^ sub-population exhibited enhanced migration in a transwell migration assay (Figure 3A). For a more stringent assessment of invasive ability, we developed a 3D matrigel-collagen invasion assay through adaptation of a previously published method (Katz et al., 2011). In this assay, the melanoma cells are embedded in a matrigel ‘plug’ that is then itself embedded within a larger collagen outer gel. In this way, the degree of invasion of melanoma cells from the matrigel plug into the surrounding collagen can be assessed. The matrigel is rich in basement membrane components, and thus provides an environment somewhat similar to the *in situ* tumour environment for a stringent assessment of invasive ability. Flattening the resulting 3D invasion into a 2D maximum intensity projection, and plotting the average distance of each invading cell from the centre of the matrigel plug, enabled assessment of the invasive ability of each sorted sub-population (Figure 3B). Both the CD24^+^CD271^+^ and CD24^+^CD271^-^ sub-populations had enhanced invasive ability compared to the CD24^-^CD271^-^ sub-population.

**Figure 3.**
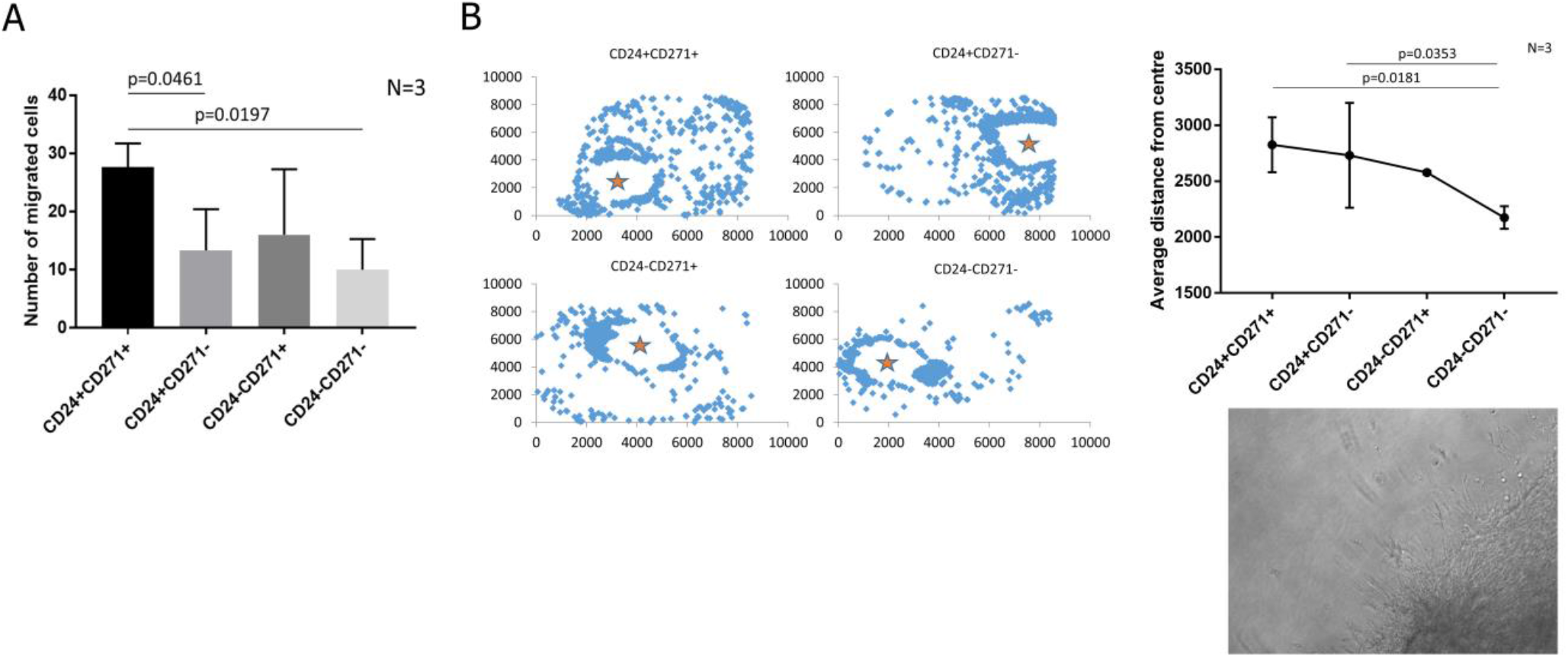
The CD24+CD271+ sub-population in the CHL-1 melanoma line has enhanced ability to migrate and invade. **A**, The number of cells that have migrated to the other side of the membrane in transwell assays, for each of the four FACS sorted sub-populations. **B**, Invasion of the four FACS sorted sub-populations into a collagen gel, when seeded in an inner matrigel gel. Left: Representative images, with each blue diamond representing a cell, projected onto x-y co-ordinates (z co-ordinates are flattened to zero). Units are pixel numbers in the x-y plane, and the orange star indicates the centre of the inner matrigel gel from which distance was calculated for each invading cell. Cells remaining in the inner gel are excluded from the visualisation. Top right: Average invasion distance for the four FACS sorted sub-populations. Bottom right: A representative brightfield image of cells breaking away from the inner gel and invading into the surrounding collagen. For each experiment, the number of biological repeats is indicated next to the graph. P-values were calculated using unpaired ANOVA.

Melanoma exhibits broad chemotherapeutic resistance, and also quickly develops resistance to targeted therapies (Kozar et al., 2019). CSCs have been reported to exhibit enhanced therapeutic resistance in many tumour types, including melanoma, and drive the development of tumour drug resistance (Biddle et al., 2016, Li et al., 2008, Phi et al., 2018). In particular, CD24^+^ CSCs in epithelial tumours exhibit the highest resistance to the chemotherapy drugs paclitaxel and cisplatin (Biddle et al., 2016), although the tyrosine kinase inhibitor dasatinib has been reported to target CD24^+^ CSCs in epithelial tumours (Goldman et al., 2015). Therefore, we assessed the relative response of the four CHL-1 sub-populations to therapeutic challenge with these drugs (Figure 4A). The CD24^+^CD271^+^ and CD24^+^CD271^-^ sub-populations became enriched after treatment with dasatinib (contrary to previous findings in epithelial tumours). Conversely, enrichment in response to paclitaxel and cisplatin was exhibited by the CD24^+^CD271^+^ and CD24^-^CD271^+^ sub-populations. Therefore, the CD24^+^CD271^-^ and CD24^-^CD271^+^ sub-populations exhibit differing resistance profiles, but these converge on the CD24^+^CD271^+^ sub-population which is enriched in response to all three drugs.

**Figure 4.**
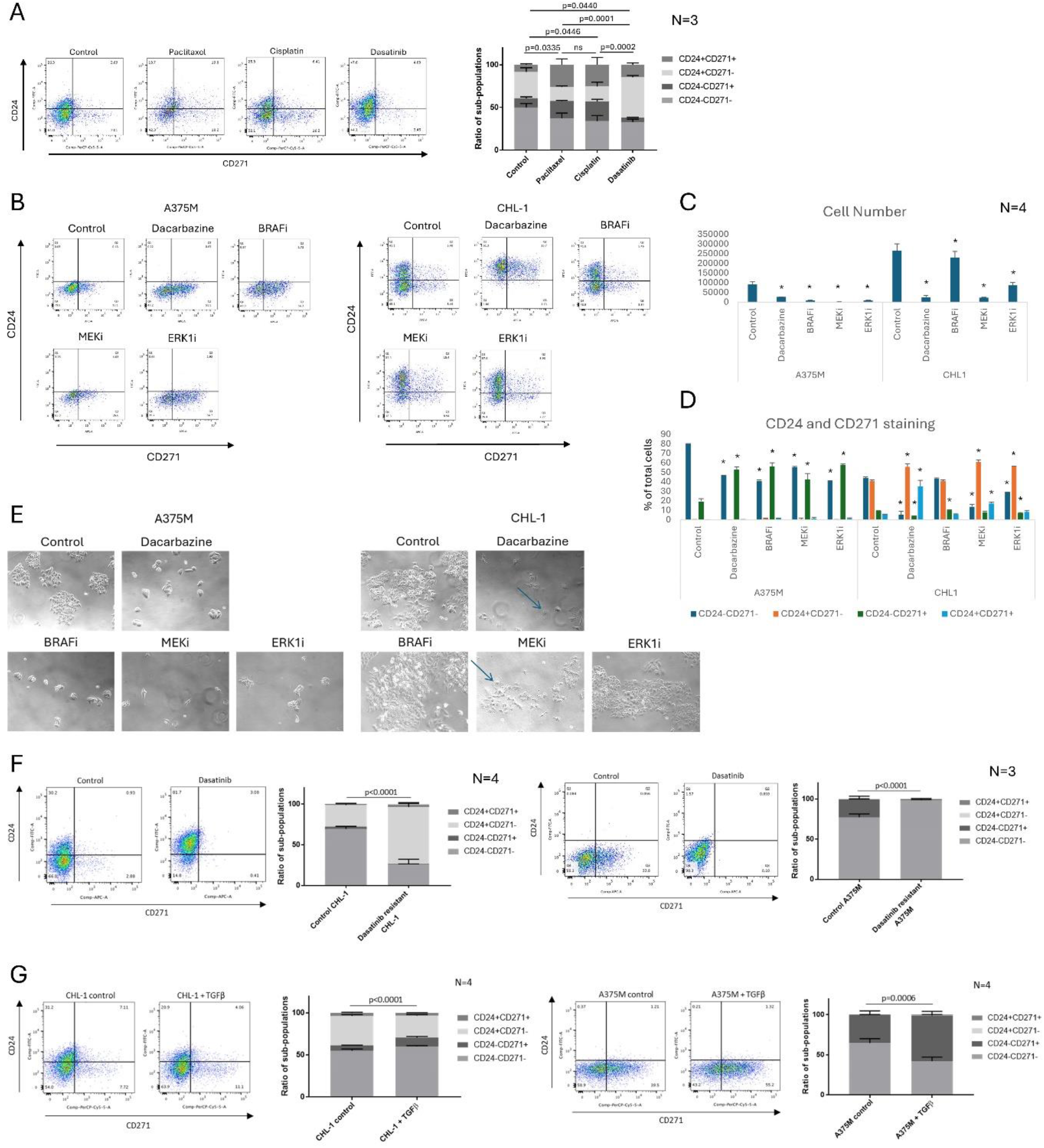
The CD24+CD271+ sub-population is drug resistant. **A**, Surviving cells in the CHL-1 melanoma line, after 3 days treatment with the indicated drugs and a 2-day recovery. **B-E**, Surviving cells in the A375M and CHL-1 melanoma lines, after 3 days treatment with the indicated drugs and a 2-day recovery. ^*^ denotes p<0.05. In the brightfield images in **E**, blue arrows indicate cells with altered morphology. **F**, Changes in distribution of sub-populations after generation of dasatinib resistant CHL-1 (left) and A375M (right) melanoma lines. **G**, Changes in distribution of sub-populations after TGFβ treatment for 6 days on CHL-1 (left) and A375M (right) melanoma lines. For all experiments, flow cytometric analysis for CD24 and CD271 is shown. The CD24/CD271 graphs show mean +/-SEM of sub-populations in each of 4 quadrants based on +/-gating set using isotype controls, as a percentage of the total cells, presented as stacked bars (**A, F, G**) or side-by-side bars (**D**). P-values were calculated using two-way ANOVA (**A, F, G**) or paired t-test (**C, D**). Representative flow cytometry plots for each condition are to the left of each graph. For each experiment, the number of biological repeats is indicated next to the graph.

We next investigated whether the presence of CD24^+^ and CD24^+^CD271^+^ sub-populations led to enhanced overall survival of the CHL-1 cell line when treated with melanoma therapeutic compounds, by comparing to the A375M cell line which contains a CD271^+^ sub-population but does not contain CD24^+^ cells. We first assessed sub-population enrichment (Figure 4B, D). The A375M cell line, which is BRAF mutant, exhibited an enrichment of the CD271^+^ sub-population in response to the chemotherapeutic agent Dacarbazine and the targeted drugs Vemurafenib (a BRAF inhibitor), Trametinib (a MEK inhibitor) and SCH772984 (an ERK1 inhibitor). The CHL-1 cell line, which is ERK1 mutant and BRAF wild type, was insensitive to BRAF inhibition. It showed an enrichment of both CD24^+^ and CD24^+^CD271^+^ sub-populations in response to Dacarbazine and Trametinib, and significant enrichment of only the CD24^+^ sub-population in response to SCH772884. Comparing overall resistance of the cell lines based on surviving cell number (Figure 4C), we observed that CHL-1 was much more proliferative than A375M under all conditions (including control) but displayed enhanced overall survival compared to A375M in response to MEK inhibition and ERK1 inhibition and no enhancement in overall survival in response to Dacarbazine. Being BRAF wild type, it was insensitive to BRAF Inhibition. Enhanced survival of CHL-1 over A375M in response to targeted agents, to which CD24^+^ characteristics appear to confer resistance, alongside no enhancement in survival to a chemotherapeutic agent, to which CD271^+^ characteristics (shared with A375M) appear to confer resistance, further highlights the differing CD24^+^ and CD271^+^ resistance profiles, and their confluence on the CD24^+^CD271^+^ sub-population.

Brightfield images of the cell lines after treatment (Figure 4E) further demonstrate the enhanced survival of CHL-1 in response to MEK inhibition and ERK1 inhibition. In concert with the cell survival data, they also show the relatively minor effect of ERK1 inhibition on the CHL-1 cell line; it continues to proliferate and display the same cellular morphology as in control conditions. Conversely, Dacarbazine and MEK inhibition induce altered morphology in the CHL-1 cell line, with the emergence of cells that exhibit a flat and spread cytoplasm (indicated by blue arrows). This effect is more pronounced with Dacarbazine, and correlates strongly with the emergence of a CD24^+^CD271^+^ sub-population (Figure 4D). This morphological effect is not observed in the A375M line, which continues to display the same cellular morphology under all control and treatment conditions.

To investigate effects on longer-term development of resistance, we treated with dasatinib, a compound that in short-term experiments targeted the CD271^+^ sub-population but simultaneously elicited enrichment of the CD24^+^ sub-populations that are absent in the A375M cell line. Long-term dasatinib treatment resulted in dasatinib resistant CHL-1 (Figure 4F, left). This caused a marked shift in the sub-population distribution, with a big increase in the CD24^+^CD271^-^ sub-population and corresponding loss of the CD24^-^CD271^+^ and CD24^-^CD271^-^ sub-populations. This underlines the role of CD24 in marking dasatinib resistant cells in this cell line. However, we were also able to create dasatinib resistant A375M using the same method, despite the absence of CD24+ sub-populations in this line. In A375M, the CD24^-^ CD271^+^ sub-population was lost, resulting in an entirely CD24^-^CD271^-^ cell line (Figure 4F, right). This stands in contrast to the previously described enrichment of the CD271^+^ sub-population upon creation of Vemurafenib resistant A375M (Patel et al., 2020). Therefore CD271^+^CD24^-^ cells, whilst resistant to BRAF inhibitor Vemurafenib, are particularly sensitive to tyrosine kinase inhibitor dasatinib. Nevertheless, the fact that we were able to create a resistant cell line from A375M, which lacks CD24+ sub-populations, indicates that long-term resistance to CSC-targeted therapy can also be induced within the CD24^-^CD271^-^ sub-population. This suggests that combination therapy, treating with CSC-targeted compounds alongside agents that kill the bulk CD24^-^CD271^-^ cell population, may be required for successful melanoma therapy.

We next considered the potential role for an amoeboid phenotype in the enhanced invasive ability of the CD24^+^ sub-populations. Amoeboid cells are known to cluster at the tumour invasive front in melanoma and drive metastatic dissemination (Georgouli et al., 2019), and this phenotype is induced by TGFβ treatment (Cantelli et al., 2015). We therefore treated the CHL-1 and A375M cell lines with TGFβ and observed the effect on the CD24 and CD271 marked sub-populations (Figure 4G). In both lines, TGFβ caused an expansion of the CD24^-^ CD271^+^ sub-population. In CHL-1, the CD24^+^CD271^-^ sub-population was reduced and there was no change in the CD24^+^CD271^+^ sub-population. Therefore, TGFβ induces the CD24^-^ CD271^+^ sub-population, indicating that an amoeboid phenotype is unlikely to be associated with the CD24^+^CD271^+^ CSC sub-population.

Finally, we investigated whether a CD24^+^CD271^+^ CSC sub-population exists in human primary melanoma specimens. FFPE sections of 31 melanomas were examined by a dermatopathologist, and included 19 superficial spreading, 11 nodular, and 1 acral lentiginous melanomas (Table 1). We developed a method for immunofluorescent co-staining for CD24 and CD271, combined with automated imaging and image stitching to create high resolution images of entire pathological block sections. This enabled accurate assessment of CD24 and CD271 co-staining in each of the specimens, which was performed by both a researcher and a pathologist, guided by the matched H&E specimen to ensure that analysis of co-staining was restricted to melanoma cells (Figure 5). Of the 31 tumours, 15 (48%) had evidence of CD24^-^CD271^+^ melanoma cells. These were specific to the invasive front in 5 (16%) tumours. Only 3 (10%) tumours had evidence of CD24^+^CD271^+^ melanoma cells and, interestingly, CD24^+^CD271^-^ melanoma cells were also restricted to these same 3 tumours. In 2 of these tumours, the staining was specific to the invasive front. These findings confirm the existence of a CD24^+^CD271^+^ sub-population that localises to the invasive front in human melanoma but, in agreement with our panel of melanoma cell lines, is only present in a minority (3/31, 10%) of tumours.

**Table 1.**
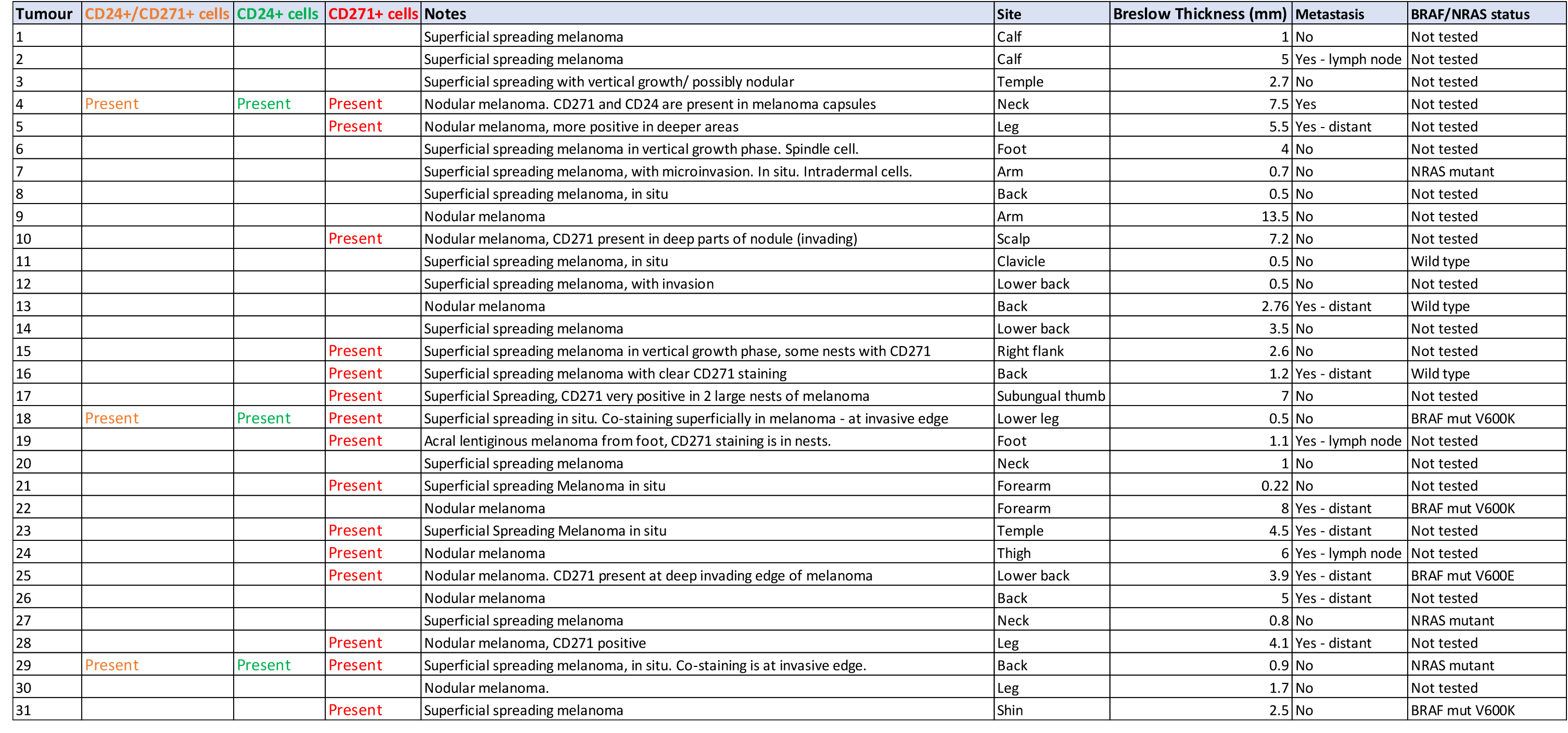
Pathological assessment of CD24 and CD271 staining in 31 melanoma specimens.

**Figure 5.**
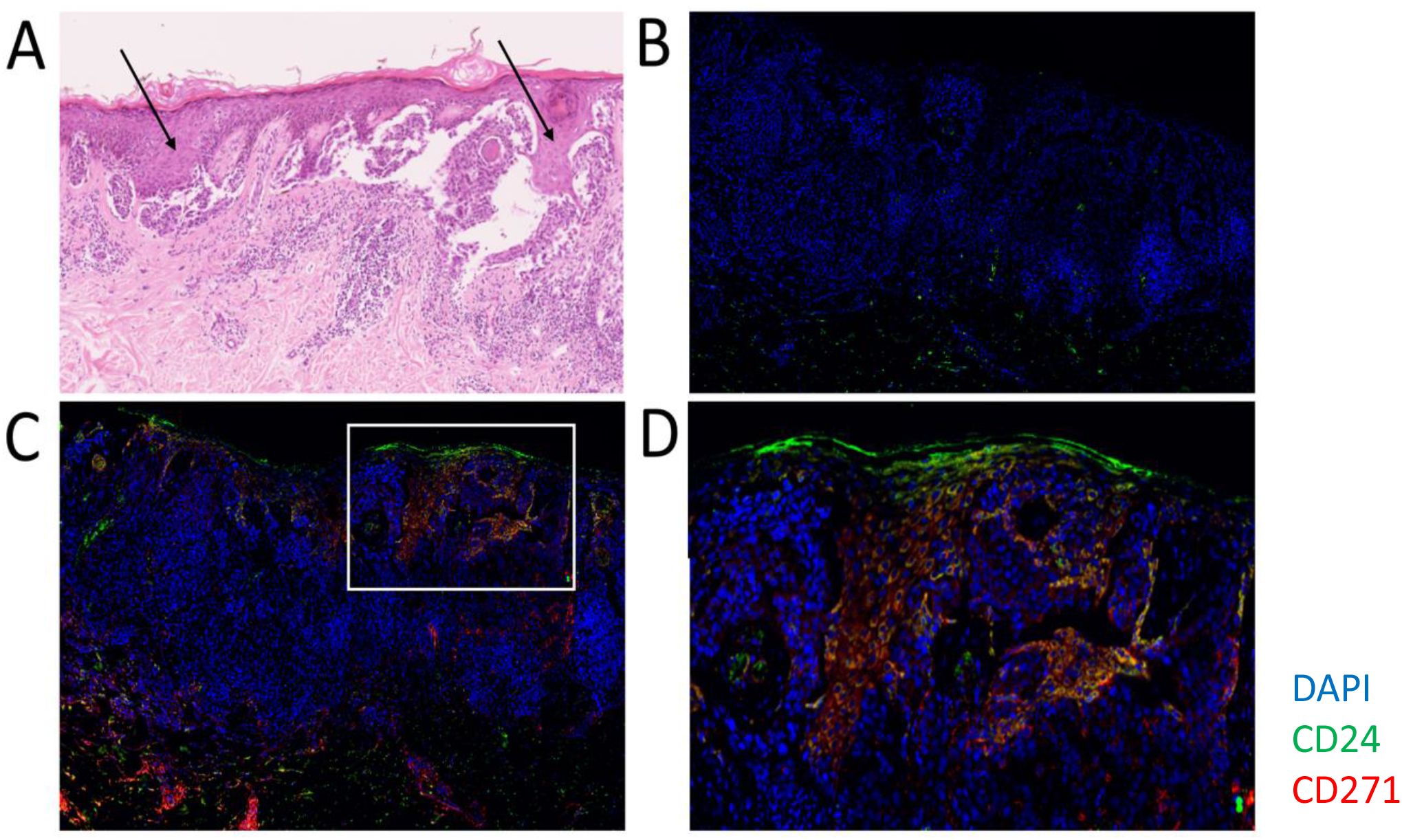
A CD24+CD271-sub-population in human melanoma tumour specimens. **A**, H&E of a human melanoma specimen, showing regions of melanoma cells (black arrows). **B**, Isotype control staining. **C**, Melanoma specimen with a region of CD24+CD271+ staining. Blue – DAPI nuclear stain, Green – CD24, Red – CD271. **D**, Magnification of the region of CD24+CD271+ staining. Displayed images are image-stitches of multiple fields of view at 40x magnification.

The occurrence of metastatic spread was strongly correlated with Breslow thickness at presentation, as is well established in melanoma (Balch et al.). It was not possible to regress out Breslow thickness in order to test association with marker staining profile within this small specimen cohort. BRAF/NRAS status was available for 10 of the specimens, and this showed no correlation between BRAF/NRAS status and CD24/CD271 staining. One of the cases containing CD24^+^CD271^+^ cells was BRAF mutant, and one was NRAS mutant. The third wasn’t tested. There were also CD24^-^CD271^+^ and CD24^-^CD271^-^ cases that were BRAF and NRAS mutant. CD24^+^CD271^+^ staining is therefore not specific to tumours that are BRAF/NRAS wildtype, nor to BRAF/NRAS mutant tumours.

## Discussion

Previous investigations of cellular heterogeneity in melanoma have led to a developing understanding of the distribution of functional sub-populations and how switching between phenotypes underlies metastasis and therapeutic resistance (Rambow et al., 2019, Müller et al., 2014). However, it is not known which (if any) melanoma cellular sub-populations exhibit CSC characteristics (Rambow et al., 2019, Quintana et al., 2010). We, and others, have previously identified a metastatic CSC sub-population that drives phenotype switching in epithelial tumours and through this propagates metastatic dissemination and therapeutic resistance (Biddle et al., 2016, Pastushenko et al., 2021, Youssef et al., 2023). In this report, we sought to test the hypothesis that a similar minority CSC sub-population exists in melanoma, whose retained lineage plasticity is responsible for driving the phenotypic switching that has been previously shown to be important for metastatic dissemination. Using the epithelial tumour CSC marker CD24 alongside the melanoma CSC marker CD271, we have identified a minority CSC sub-population that exhibits enhanced self-renewal and lineage plasticity. It is also more invasive and exhibits heightened drug resistance. This CD24^+^CD271^+^ CSC sub-population exhibited a confluence of the characteristics of discrete CD24^+^ and CD271^+^ sub-populations. Unlike the CD271^+^ sub-population, the CD24^+^CD271^+^ and CD24^+^ sub-populations only occurred in 1 out of 4 established melanoma cell lines and 3 out of 31 melanoma tumour specimens. There was no association with BRAF/NRAS mutation status.

Looking at mutation status in the cell lines used in this study; LM4 is BRAF mutant and NRAS wild-type, LM32 is wild-type for both (Daniotti et al., 2004), CHL-1 is wild-type for both, and A375M is BRAF mutant and NRAS wild-type (ATCC). Therefore, as with the tumour specimens, there is no correlation between BRAF/NRAS mutation status and presence of CD24^+^ sub-populations in the cell lines. The CHL-1 line has an ERK1 mutation, and yet displays substantial resistance and CD24^+^ sub-population enrichment in response to ERK1 inhibition. Therefore, components of the BRAF/NRAS/ERK1 pathway do not appear to be responsible for the observed therapeutic resistance of the CHL-1 cell line and its CD24^+^ and CD24^+^CD271^+^ sub-populations. Interrogation of the TGGA melanoma cohort (cBioPortal) supports this conclusion, showing no correlation of CD24 expression with BRAF, NRAS or ERK1 mutation or expression (though only 2% of cases are ERK1 mutant). The Broad Institute melanoma immunotherapy scRNAseq dataset (Jerby-Arnon et al., 2018) has one tumour out of 14 containing CD24^+^ cells. This was in a bi-modal distribution, with only some of the cells being CD24^+^. There was no correlation between CD24 and BRAF or ERK1 expression in these cells. Interestingly, whilst NRAS was expressed over a continuous distribution in both the CD24^+^ and CD24^-^ sub-populations in this tumour, the two sub-populations differed in that most of the CD24^+^ cells were towards the high end of the NRAS expression distribution. This suggests a potential role for high NRAS expression, but not mutant NRAS, in the CD24^+^ sub-population. Interestingly, in our experiments, inhibition of the NRAS effector MEK caused greater cell death and CD24^+^CD271^+^ population induction than BRAF or ERK1 inhibition in the CHL-1 cell line. Perhaps, as with the chemotherapeutic drug Dacarbazine, confluence of CD24^+^ and CD271^+^ attributes is required for optimal resistance to MEK inhibitors.

QPCR suggested a neural crest-like state of the CD24^+^ cells, but with heightened proliferation and reduced Notch signalling. Notch signalling is increased in slow-cycling EMT-like and CD271^+^ cells in melanoma (Murtas et al., 2017, Filipp et al., 2019), so the reduced Notch signalling in the CD24^+^ cells suggests they are an alternative, non-EMT-like state. TGFβ is an inducer of the EMT-like as well as amoeboid-like state in melanoma (Cantelli et al., 2015), so reduction of the CD24^+^ and increase in the CD271^+^ populations in response to TGFβ further supports the non-EMT-like status of the CD24^+^ cells. These findings suggest that the CD24^+^ cells represent an alternative, proliferative, neural crest-like state that stands apart from the previously described slow-cycling EMT-like CD271^+^ neural crest-like state (Caramel et al., 2013).

Dasatinib treatment induced almost total loss of the CD271^+^ sub-population, alongside a big increase in the CD24^+^ sub-population. In unpublished work on OSCC cell lines (L. Gammon, unpublished), Dasatinib selectively removed cells that have undergone EMT. Therefore, in melanoma, there may be a parallel process where Dasatinib removes CD271^+^ EMT-like neural crest-like cells whilst simultaneously selecting for CD24^+^ non-EMT-like neural crest-like cells. A key target of Dasatinib is Src tyrosine kinase, and there is evidence that Src connects Notch signalling to downstream HIF1α signalling (Lee et al., 2009), which is itself an inducer of EMT (Yang et al., 2008). Therefore, Dasatinib may sever this link to inhibit the production of an CD271^+^ EMT-like state and instead favour production of CD24^+^ non-EMT-like neural crest-like cells.

Previous findings have shown that CD24 expression in primary melanoma is one of the most predictive markers for metastatic disease (Gschaider et al., 2012). Similarly, Tang et al. (2014) demonstrated that CD24 was increased in melanoma patient samples compared to normal tissue, and that this marker was prognostic, with elevated CD24 correlating with decreased survival (Tang et al., 2014). A pro-metastatic CD24+ sub-population has also been characterised in a mouse model of melanoma (Liu et al., 2012). Therefore, whilst CD24+ sub-populations were only seen in a minority of tumours, they may confer characteristics that are important for metastatic spread.

The CD24^+^CD271^+^ sub-population was resistant to all tested drug treatments, and appeared to combine the resistance profiles of the separate CD24^+^ and CD271^+^ sub-populations into a hybrid profile with broad therapeutic resistance. We observed the emergence of an altered morphology in the CHL-1 cell line, that correlated with the emergence of a CD24^+^CD271^+^ sub-population upon drug treatment. The cells exhibited a flat and spread cytoplasm, similar to that seen in a recent study where the emergence of ‘giant cells’ was observed in response to chemotherapeutic treatment of a mouse melanoma cell line (Weng et al., 2020). In this previous study, giant cells were proliferative, chemo-resistant, and expressed melanoma stem cell markers ABCB5 and CD133. Cytoplasmic spreading has also been previously seen in response to CD24 expression, which was associated with enhanced proliferation, invasion and metastatic spread in breast cancer (Baumann et al., 2005). This, alongside other CD24^+^CD271^+^-specific attribute including tumoursphere formation, may be an important feature of this CSC sub-population.

Interestingly, in epithelial tumours, tumoursphere formation in serum-containing FAD medium is restricted to sub-populations that have undergone EMT (Biddle et al., 2011, Biddle et al., 2016). The mesenchymal character of the melanoma cell lines maybe explains their ubiquitous sphere-forming abilities under these conditions. Conversely, under more stringent serum-free conditions, sphere-forming ability was restricted to the CHL-1 cell line that contains the CD24^+^CD271^+^ sub-population and was heightened in this sub-population. Given that sphere-forming cells must resist oxidative stress, this draws parallels with work demonstrating that metastasising melanoma cells must resist oxidative stress in the circulation and undergo reversible metabolic changes to achieve this (Piskounova et al., 2015). Therefore, the CD24^+^CD271^+^ cells may represent a minority CSC sub-population that is primed for efficient metastatic dissemination. Thus, functional requirement for CD24^+^CD271^+^ CSCs may only manifest under particular conditions, for example the increased oxidative stress associated with metastatic dissemination or therapeutic challenge. However, even here there may be other ways for melanoma to subvert the requirement for this CSC sub-population, for example by metastasising via the lymphatics where they experience less oxidative stress (Ubellacker et al., 2020). Alongside the long-term drug resistance findings described here, where even CD24^-^CD271^-^ cells can support the emergence of resistance to targeted therapies, this underlines the potential importance of developing combination therapies to simultaneously target all cellular sub-population in melanoma.

## Methods

### Cell culture

Four human melanoma lines were used in this study: CHL-1 (provided by Penny Lovat, Newcastle), A375M (provided by John Marshall, Barts & the London), LM4 and LM32 (provided by Monica Rodolfo, Inst. Tumori Milano). LM4 was from a human primary tumour of the trunk region (Daniotti et al., 2004). CHL-1 (RRID: CVCL_1122) and LM32 (Daniotti et al., 2004) were from human metastatic sites (pleural effusion and cutaneous metastasis, respectively). A375M was formed from pooled lung metastatic deposits of a human melanoma cell line in a mouse metastatic model (Kozlowski et al., 1984). Cell culture was performed as previously described (Biddle et al., 2011). Cell removal from adherent culture was performed using 1x Trypsin-EDTA (Sigma, T3924) at 37°C.

### Flow cytometry and fluorescence-activated cell sorting (FACS)

Flow cytometry was performed as previously described (Biddle et al., 2011). Antibodies for cell line staining were CD44-PE (clone G44-26, BD Bioscience), CD24-FITC (clone ML5, BD Bioscience, and clone REA832, Miltenyi Biotec), CD271-PerCP/Cy5.5 and CD271-APC (both clone C40-1457, BD Bioscience), and EpCAM-APC (clone HEA-125, Miltenyi Biotec). Single stained controls were performed for compensation, and isotype controls were performed to set negative gating. Flow cytometry data was analysed using FlowJo software, and statistical testing performed using 2-way ANOVA with multiple comparisons in GraphPad Prism software (unless otherwise indicated in figure legend).

### Sphere assay

For suspension cultures, 48-well plate wells were coated with polyhema (Sigma) (12mg/ml in 95% ethanol). Cells were then plated at a density of 1000 cells/ml, incubated (37C, 5% CO2), and spheres were counted after 7 days. 1x Trypsin-EDTA was used to break down spheres for secondary sphere plating. The medium for suspension culture was either fully supplemented FAD medium (Biddle et al., 2011) or defined serum free medium (Sato et al., 2011), as indicated, with addition of 1% methylcellulose (Sigma). Statistical testing was performed using paired ANOVA with multiple comparisons.

### QPCR

RNA extraction, cDNA synthesis and QPCR were performed as previously described (Biddle et al., 2011). Primer sequences were; Zeb1 F: GTCCAAGAACCACCCTTGAA R: TTTTTGGGCGGTGTAGAATC; Vimentin F: CCCTCACCTGTGAAGTGGAT R: GACGAGCCATTTCCTCCTTC; Axl F: CAAGTGGATTGCCATTGAGA R: GATATGGGGTTTGGCCTCTT; Cyclin D1 F: CTTCCTCTCCAAAATGCCAG R: TGAGGCGGTAGTAGGACAGG; MITF F: GGGCTTGATGGATCCTGCTT R: TATGTTGGGAAGGTTGGCTG; Hes1 F: TGAGCCAGCTGAAAACACTGA R: TGCCGCGAGCTATCTTTCTT; Hey1 F: TAATTGAGAAGCGCCGACGA R: GCTTAGCAGATCCCTGCTTCT GAPDH F: GTGAACCATGAGAAGTATGACAAC R: CATGAGTCCTTCCACGATACC.

### Colony assay

Cells were plated at a concentration of 1,000 cells per ml and cultured for seven days. They were then fixed in 4% paraformaldehyde and stained with crystal violet for colony counting. Statistical testing was performed using unpaired ANOVA with multiple comparisons.

### Migration assay

1000 cells were plated in Transwell tissue culture inserts (8 mm membrane, Corning) in 24-well plates. After 24 hours, the membranes were fixed in 4% paraformaldehyde, stained with crystal violet, the non-migrated cells on the top of the membrane removed with a cotton-wool bud, and the migrated cells on the underside of the membrane counted. Statistical testing was performed using unpaired ANOVA with multiple comparisons.

### Matrigel-Collagen invasion assay

10,000 cells were suspended in 50µl matrigel (Corning) in 96-well plate wells, which was allowed to set before being removed and inserted into the middle of 500µl 4mg/ml collagen I (Corning) in 48-well plate wells coated with polyhema. Once this had set, medium was added and the cells were cultured for 7 days. For the final day of culture, 20 µg/ml Hoescht 33342 (Sigma) was added to the medium. The cultures were then fixed in 4% paraformaldehyde, and imaged through z-stacks using an InCell 2200 automated epifluorescent microscope (GE) to detect all stained nuclei. For each stained nucleus, distance from the centre of the small inner gel in the x-y plane was calculated and reconstructed in a graphical plot. Average distance was calculated for each sorted sub-population, and statistical testing was performed using unpaired ANOVA with multiple comparisons.

### Drug treatment, generation of dasatinib resistant lines, and TGFβ treatment

Cells were plated at a density of 4000 cells per ml in T25 flasks. Drugs were added to the following final concentrations: Cisplatin 1.5 μM (Sigma), Paclitaxel 10 μM (Sigma), Dasatinib 0.5 μM (BioVision), Dacarbazine 1 mM (Abcam), Vemurafenib 2 μM (Bio-Techne), Trametinib 50 nM (Bio-Techne), SCH772984 100 nM (Cayman). Medium was replaced after 3 days to remove the drugs and cells were analysed by flow cytometry after a further 2 day recovery period. For the generation of Dasatinib resistant CHL-1 and A375M, cells were grown in 1 µM Dasatinib continuously with the medium and drug changed every 3 days until robust growth was re-established. Cells were then analysed by flow cytometry. For TGF-β treatment, 2 ng/ml TGF-β (R&D Systems) was added. TGF-β was replaced once during a six days incubation period, followed by flow cytometric analysis.

### Ethical approval for human melanoma samples

This study was approved by the East London Research Ethics Committee (project 07/Q0604/23). Consent was obtained from all patients in accordance with the Declaration of Helsinki.

### Immunofluorescent staining of tumour tissue sections

FFPE blocks from were obtained from Barts NHS Trust histopathology archive. H&E slides were examined by a dermatopathologist to confirm the diagnosis of melanoma and specific histological features (table 1). 5 µm sections on poly-lysine slides were then de-waxed in xylene and ethanol and hot antigen retrieval was performed in Tris-EDTA pH9 for 10 minutes. Permeabilisation was performed for 10 minutes in 0.25% Triton X-100 in PBS. Washes were in PBS, and blocking was overnight at 4 degrees C with 3% BSA in PBS. Primary antibody incubation was overnight at 4 degrees C with the following antibodies at 1:100 dilution in PBS + 3% BSA: CD24-FITC (clone ML5, BD Bioscience), CD271-APC (clone C40-1457, BD Bioscience). Isotype controls were included. After washing in PBS + 3% BSA, sections were incubated for 1 hr at room temp in the following secondary antibodies at 1:500 dilution in PBS + 3% BSA: Alexa Fluor 488 Goat Anti-mouse IgG2a (Invitrogen) (binds the CD24 primary antibody), Alexa Fluor 647 Goat Anti-mouse IgG1 (Invitrogen) (binds the CD271 primary antibody). Antibody combinations were tested for cross reactivity, and none was seen. After washing again in PBS + 3% BSA, sections were incubated in 1 µg/ml DAPI (Sigma) for 10 minutes, washed in PBS, and mounted using Immu-mount (Shandon). Immunofluorescent imaging was performed using an INCA2200 Analyser (GE), capturing between 80 and 108 overlapping 40x fields in a region with confirmed melanoma as judged by pathological assessment of the H&E. This was performed in three blocks per sample. Images were stitched into blocks of 36 fields of view for further analysis.

### Identifying sub-populations in tumour tissue sections

Stitched images were examined by a researcher and a dermatopathologist to identify areas of positive, true (non-background) CD24+ and CD271+ staining, and these were compared to the corresponding area of the H&E. Areas of single staining (CD24+ or CD271+) and double staining (CD24+/CD271+) on the stained images were recorded if they coincided with melanoma cells. The nature and location of this staining was also recorded.

## Conflict of interest

The authors declare no conflicts of interest.

### Acknowledgements

We thank Gary Warnes for technical assistance and discussion. Cell lines were kindly provided by Penny Lovat, Newcastle; John Marshall, Barts & the London; Monica Rodolfo, Inst. Tumori Milano. Adrian Biddle has received funding support from Animal Free Research UK, as part of the Animal Replacement Centre of Excellence at Queen Mary University of London, and from the UK Medical Research Council (MR/V009494/1). Adrian Biddle is a member of the Barts Centre for Squamous Cancer, funded by Barts Charity (G-002030).

## CRediT statement

Conceptualisation: AB; Methodology: OK, PD, IH-R, SL, GY, LG; Formal analysis: OK, HR; Investigation: OK, PD, IH-R, HR, AB; Resources: ICM, MPP, DB, CAH; Data curation: CAH; Writing – original draft: AB; Writing – review and editing: OK, PD, CAH; Visualisation: OK, PD, IH-R, AB; Supervision: ICM, MPP, AB.

